# Single-Cell Sequencing in Pediatric Acute Leukemia: A Systematic Review of Biologic Insights and Translational Potential

**DOI:** 10.64898/2026.10.06.757108

**Authors:** Jacopo Martelli

**Author notes:** Correspondence: Jacopo Martelli.

## Abstract

**Background:** Relapse remains the principal cause of treatment failure in pediatric acute leukemia. Clinically validated minimal residual disease (MRD) assays — multiparameter flow cytometry, immunoglobulin/T-cell-receptor rearrangement sequencing and targeted next-generation sequencing panels — are highly sensitive but read out predefined features and do not reconstruct clonal architecture or link a cell’s genotype to its functional state. We systematically reviewed what single-cell sequencing of pediatric acute leukemia has revealed about clonal evolution, chemoresistance, the immune microenvironment and post-therapeutic immune dynamics.

**Methods:** This review followed PRISMA 2020. PubMed/MEDLINE, the Cochrane Library and Google Scholar were searched to September 2026 for primary studies applying single-cell DNA, RNA, chromatin-accessibility, surface-protein or immune-repertoire sequencing to pediatric acute leukemia samples or pediatric-derived models. Risk of bias in non-randomized studies was assessed with ROBINS-I. Because designs, platforms and outcomes were heterogeneous, findings were synthesized narratively by biologic theme without meta-analysis, following the SWiM guideline.

**Results:** Of 63 records identified, 10 studies were included (published 2016–2026; 6–159 patients or samples per study; up to *∼*700,000 cells profiled). Single-cell DNA sequencing traced minor *NOTCH1*-mutant diagnostic subclones into *PTEN*-or *TP53*-mutant relapse clones that bulk sequencing missed. Rare dormant, label-retaining cells and developmental-state-dependent drug sensitivity defined chemoresistant compartments. Remodeling of the bone marrow immune microenvironment identified patients with adverse outcome among those classified as MRD-low, and pre-infusion CAR T-cell type 2 functionality and post-transplant cytotoxic memory CD8^+^ T-cell subsets were associated with durable remission. Of the seven studies assessed with ROBINS-I, four were at moderate and three at serious overall risk of bias, driven chiefly by small, selected cohorts.

**Conclusions:** Single-cell sequencing yields mechanistic insight that complements, rather than replaces, clinically validated MRD assays, and nominates candidate biomarkers for risk stratification, treatment selection, cell-therapy manufacturing and post-transplant monitoring. The evidence remains preliminary; adequately powered, prospective and harmonized studies are required before clinical adoption.

**Registration:** This review was not prospectively registered.

## 1 Introduction

### 1.1 Clinical context

Acute lymphoblastic leukemia (ALL) and acute myeloid leukemia (AML) are the most common malignancies of childhood. Contemporary protocols achieve 5-year event-free survival of approximately 85–90% [1], and 5-year overall survival now exceeds 90% for childhood ALL in high-income regions [2]. Relapse nevertheless occurs in 15–20% of patients and carries a markedly worse prognosis, with post-relapse overall survival of approximately 50% for BALL, 35% for T-ALL and 21% for infant ALL (Figure 1) [3]. Minimal residual disease (MRD) — leukemic cells persisting below the detection threshold of morphologic assessment — is the most powerful independent predictor of relapse [4].

**Figure 1:**
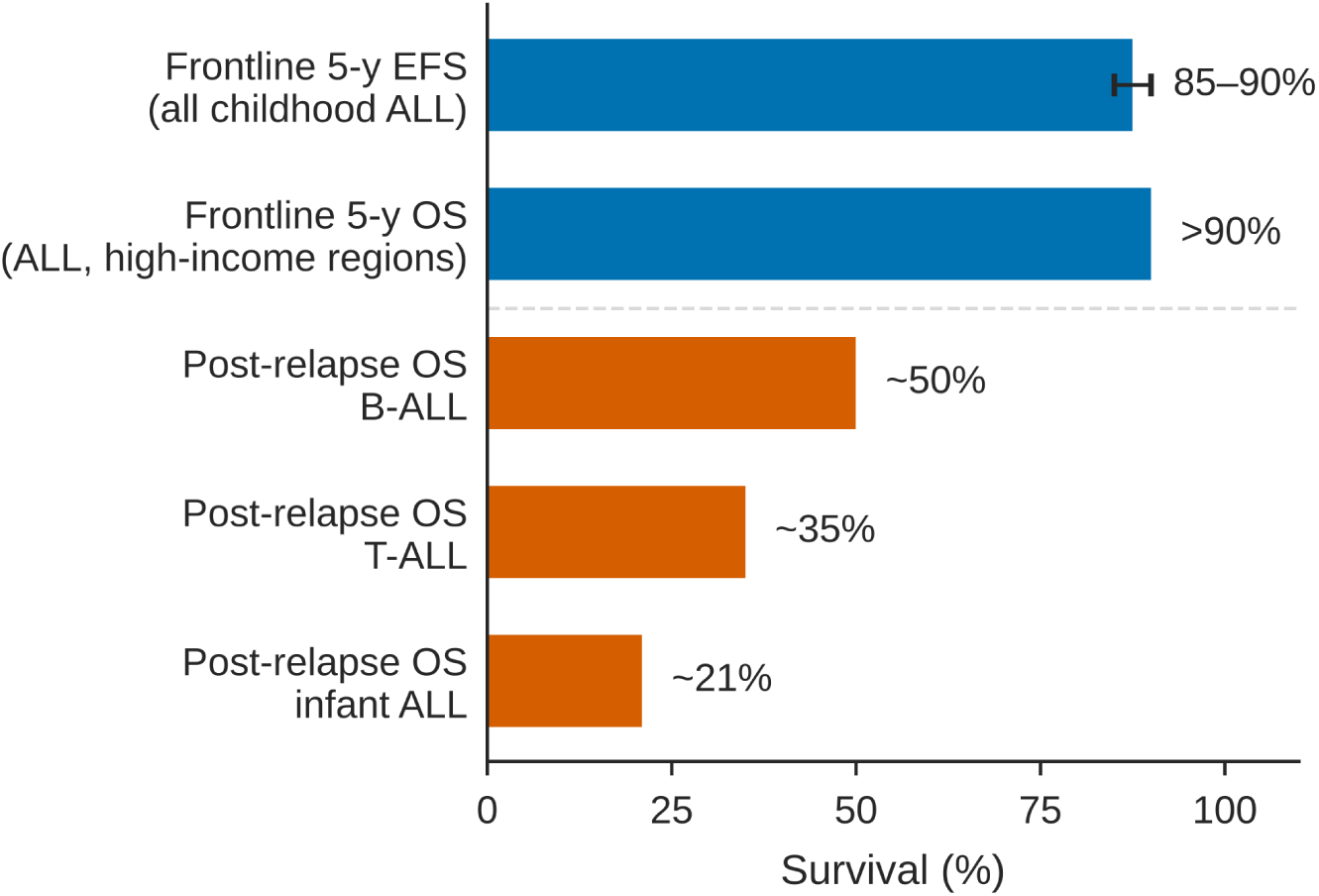
Survival of childhood acute lymphoblastic leukemia at first diagnosis versus after first relapse. Five-year event-free survival (EFS) with contemporary frontline protocols (range 85–90%) [1] and five-year overall survival (OS) in high-income regions [2] contrasted with OS after first relapse by subtype [3]. Values are approximate summary estimates from the cited sources, not pooled data.

Clinical MRD monitoring relies on multiparameter flow cytometry or molecular assays, including high-throughput sequencing of immunoglobulin and T-cell-receptor gene rearrangements such as the clonoSEQ assay [5]; targeted next-generation sequencing (NGS) gene panels additionally inform risk assessment and the selection of targeted therapy [6]. These assays are clinically validated, highly sensitive and, in the case of rearrangement sequencing, explicitly clone-aware. They are not designed, however, to reconstruct within a single assay the full clonal architecture of a leukemic population — the relative abundance, co-occurring mutations and lineage relationships of every subpopulation — or to link a cell’s genotype to its transcriptional and differentiation state. Consequently, they may under-represent subclones below their sensitivity floor and give limited visibility into the evolutionary changes that precede relapse, such as the emergence of subclones with new driver mutations, shifts in clonal dominance and changes in differentiation state [7].

### 1.2 Single-cell sequencing technologies

Single-cell sequencing profiles the genome, transcriptome, chromatin accessibility or surface proteome of thousands of individual cells in parallel, resolving the cellular heterogeneity that underlies treatment resistance and relapse [8]. All droplet-based methods share one principle: each cell is encapsulated in its own aqueous droplet, and the molecules released from it are tagged with a droplet-specific barcode, so that every sequencing read can be traced back to its cell of origin and a cell *×* feature matrix can be reconstructed from one experiment (Figure 2A). The modalities differ in which layer of cellular biology they read out (Figure 2B; Table 1). For clinicians without a computational background, the proliferation of platform names can obscure what is essentially an intuitive idea: each technology measures one biological layer one cell at a time, rather than as an average across a population.

**Figure 2:**
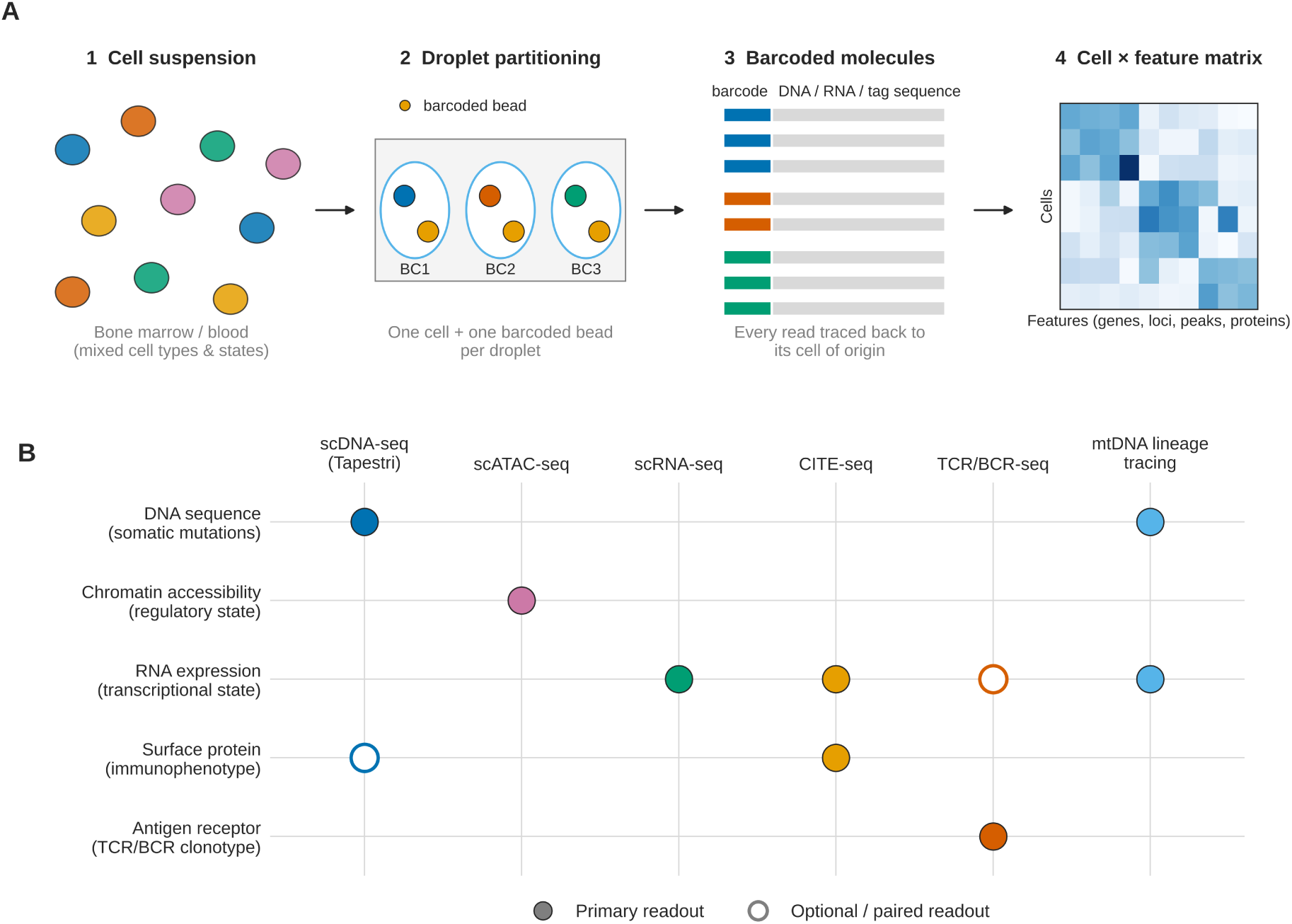
Principle of droplet-based single-cell sequencing and the biological layer measured by each modality. (A) Shared workflow: a dissociated bone marrow or blood sample is partitioned so that each droplet contains one cell and one barcoded bead; all molecules from that cell receive the same barcode, so every read can be assigned to its cell of origin, yielding a cell *×* feature matrix. (B) Modalities (columns) mapped to the layer of cellular biology they read out (rows). Filled circles, primary readout; open circles, optional or paired readout (protein tags on Tapestri; transcriptome paired with TCR/BCR sequencing). Mitochondrial-DNA lineage tracing recovers mtDNA variants and gene expression from the same cell [9, 10].

**Table 1:** Single-cell sequencing modalities discussed in this review.

| **Modality** | **What it measures** | **Core technical principle** | **Included studies using it** |
| --- | --- | --- | --- |
| Single-cell DNA sequencing | Somatic mutations and clonal genotype per cell | Droplet encapsulation; targeted amplicon PCR of a gene panel; reads barcoded per cell | Albertí-Servera et al. [11] |
| scRNA-seq | Gene expression / transcriptional state | Barcoding of mRNA from each cell during reverse transcription | Nine of ten studies (Table 3) |
| scATAC-seq | Chromatin accessibility / regulatory state | Tn5 transposase inserts adapters preferentially into open chromatin of each nucleus | Liu [13]; Bai [20]; Li [21]; Shoeb [22] |
| CITE-seq | Surface-protein immunophenotype with transcriptome | DNA-barcoded antibodies sequenced alongside the transcriptome of the same cell | Wiggers [16]; Bai [20] |
| TCR/BCR repertoire sequencing | Clonal identity of individual T and B cells | Sequencing of the hypervariable CDR3 receptor region | Wiggers [16]; Suo [23] |
| Computational lineage tracing | Clonal evolutionary relationships over time | Reconstruction of clonal ‘family trees’ from shared somatic (nuclear or mitochondrial) mutations across cells | Albertí-Servera [11]; Liu [13] (clonal/state inference) |

#### Single-cell DNA sequencing

Platforms such as Tapestri lyse each encapsulated cell and perform targeted PCR amplification of a panel of recurrently mutated genes (for example, 305 amplicons across 110 leukemia genes) [11]. Because amplicons are barcoded per cell, the platform reports which combination of mutations each cell carries — the information required to reconstruct clonal architecture — and oligonucleotide-conjugated antibodies can add surface-protein measurements from the same cells. Bulk sequencing, in contrast, can report only that mutations are present somewhere in the sample. The main technical limitation is allele dropout, in which one allele fails to amplify from the minute quantity of single-cell DNA.

#### Single-cell RNA sequencing (scRNA-seq)

Messenger RNA from each cell is captured and barcoded during reverse transcription, producing a cell *×* gene expression matrix. Because expression reflects functional state, scRNA-seq is the primary tool for distinguishing cell types and states — dormant stem-like cells, cycling blasts, mature immune cells — and for identifying expression signatures associated with outcomes such as MRD status [12]. Its principal limitation is transcript dropout.

#### Single-cell chromatin accessibility (scATAC-seq)

The Tn5 transposase preferentially inserts sequencing adapters into open chromatin, so accessibility can be read out per nucleus. Because chromatin remodeling often precedes changes in expression, scATAC-seq can reveal regulatory programs — including shifts toward stem-like or resistant states — that are not yet apparent transcriptionally, and has distinguished chemoresistant from chemosensitive subclones with superficially similar transcriptomes [13, 14].

#### CITE-seq and immune-repertoire sequencing

CITE-seq measures surface proteins with DNA-barcoded antibodies alongside the transcriptome in the same cell, without the spectral-overlap limit of fluorescent flow cytometry [15]. It is frequently combined with T-and B-cell receptor sequencing, in which the hypervariable CDR3 region serves as a natural clonal fingerprint of each lymphocyte [16].

#### Lineage tracing

Engineered, CRISPR-edited barcodes can record cell-division histories but require genetic manipulation and are restricted to experimental systems [17]. In primary human samples, somatic mitochondrial DNA (mtDNA) variants serve as endogenous, high-copy barcodes inherited by daughter cells [9]; enrichment of mitochondrial transcripts during scRNA-seq library preparation recovers clonal identity and expression state from the same cell [10]. Nuclear driver mutations such as *NOTCH1* are present in only two copies per cell and subject to dropout, making them unreliable as stand-alone lineage barcodes, although their cooccurrence patterns across many cells remain highly informative for clonal reconstruction. The resulting ability to detect rare chemoresistant subpopulations and trace them from diagnosis to relapse is the central biologic contribution of these technologies [18, 19].

### 1.3 Rationale and objectives

Single-cell studies of pediatric acute leukemia have multiplied rapidly, but they span heterogeneous platforms, disease subtypes and study designs, and their findings are largely inaccessible to clinicians without a computational background. This systematic review aimed to identify and synthesize primary studies applying single-cell sequencing to pediatric acute leukemia in order to answer four questions: (i) how leukemic clones evolve during treatment; (ii) which cell states underlie chemoresistance; (iii) how the immune microenvironment relates to MRD and outcome; and (iv) which immune cell states after CAR T-cell therapy or hematopoietic stem cell transplantation (HSCT) predict durable remission. Secondary aims were to appraise the risk of bias of this evidence and to identify the barriers to clinical translation.

## 2 Methods

### 2.1 Protocol, registration and reporting

This review was conducted and reported in accordance with the Preferred Reporting Items for Systematic Reviews and Meta-Analyses (PRISMA) 2020 statement [24]; the completed checklist is provided in Supplementary Table S3. Because no quantitative pooling was performed, the synthesis is reported following the Synthesis Without Meta-analysis (SWiM) guideline [25]. A review protocol was not prospectively registered.

### 2.2 Eligibility criteria

Eligible studies were primary research articles that (i) applied single-cell DNA, RNA, chromatin-accessibility, surface-protein or immune-repertoire sequencing, with or without lineage tracing; (ii) to samples from patients with pediatric acute leukemia (ALL, AML, mixed-phenotype acute leukemia, acute megakaryoblastic leukemia or juvenile myelomonocytic leukemia; age *≤*18 years at diagnosis), to mixed-age cohorts that included pediatric patients, or to experimental models derived from or specific to pediatric acute leukemia (patient-derived xenografts or engineered stem-cell models of pediatric-specific fusions); and (iii) reported findings on clonal evolution, chemoresistance, the immune microenvironment, MRD or post-therapeutic immune dynamics. Studies restricted to adult cohorts, non-leukemic malignancies, reviews, editorials or commentaries without primary data, and non-English-language publications were excluded.

### 2.3 Information sources and search strategy

PubMed/MEDLINE, the Cochrane Library and Google Scholar were searched from inception to September 2026 using Medical Subject Headings (MeSH) and title/abstract keywords combined with Boolean operators (Table 2). The full query-by-query PubMed yield is reported in Supplementary Table S1.

**Table 2:** Search terms applied (PubMed/MEDLINE syntax; adapted for the Cochrane Library and Google Scholar).

| Search no. | Search terms |
| --- | --- |
| #1 | leukemia [MeSH Terms] |
| #2 | analysis, single cell [MeSH Terms] |
| #3 | multiomics [MeSH Terms] |
| #4 | minimal residual disease [MeSH Terms] |
| #5 | child [MeSH Terms] |
| #6 | #1 AND #2 AND #4 AND #5 |
| #7 | #1 AND #3 AND #4 |
| #8 | scRNA-seq[Title/Abstract] OR scDNA-seq[Title/Abstract] OR scATAC-seq[Title/Abstract] |
| #9 | #1 AND #4 AND #8 |
| #10 | single cell multi omics[Title/Abstract] |
| #11 | resistant[Title/Abstract] |
| #12 | #1 AND #10 |
| #13 | #1 AND #10 AND #11 |
| #14 | Leukemia [All Fields] |
| #15 | Pediatric OR Child OR Adolescent [All Fields] |
| #16 | Single cell multi omics [All Fields] |
| #17 | resistant clones OR minimal residual disease [All Fields] |
| #18 | #14 AND #15 AND #16 AND #17 |

### 2.4 Selection process

Records were imported into Rayyan [26] for deduplication. Titles and abstracts were screened by J.M., and potentially eligible records were assessed in full text against the eligibility criteria.

### 2.5 Data collection and data items

From each included study the following items were extracted: first author, year, country, study design, population or model system, sample size, single-cell modalities, number of cells profiled, principal findings and authors’ or reviewers’ noted limitations. Data were extracted by J.M.

### 2.6 Risk-of-bias assessment

Risk of bias was assessed with the Risk Of Bias In Non-randomised Studies of Interventions (ROBINS-I) tool across seven domains: confounding, selection of participants, classification of interventions, deviations from intended interventions, missing data, measurement of outcomes and selection of the reported result [27]. Each domain was judged as low, moderate, serious or critical risk, and the overall judgement was set to the most severe domain-level judgement, as specified by the tool. ROBINS-I was applied to the seven studies with an observational or model-based design amenable to domain-level appraisal; it was not applied to the two predominantly xenograft-based experimental studies [28, 29] or to the diagnostic-accuracy study [23], for which the tool is not designed, and their limitations are described qualitatively.

### 2.7 Synthesis methods

Marked heterogeneity in design, single-cell platform, disease subtype and outcome definitions precluded meta-analysis. Studies were therefore grouped according to the principal biologic question they addressed — clonal evolution, chemoresistance, the immune microenvironment, and post-therapeutic immune dynamics — and synthesized narratively within each theme; studies informing more than one theme are discussed in each. Neither effect measures nor vote counting were used. Relevant single-cell studies identified outside the structured search are cited separately as contextual evidence, are explicitly labelled as such, and were not included in the formal synthesis or risk-of-bias assessment. Reporting biases and the certainty of the body of evidence were not formally assessed (see Section 4.4).

## 3 Results

### 3.1 Study selection

The search identified 63 records. After removal of 2 duplicates, 61 records were screened by title and abstract and 51 were excluded. All 10 remaining reports were assessed in full text, met the eligibility criteria, and were included in the synthesis (Figure 3).

**Figure 3:**
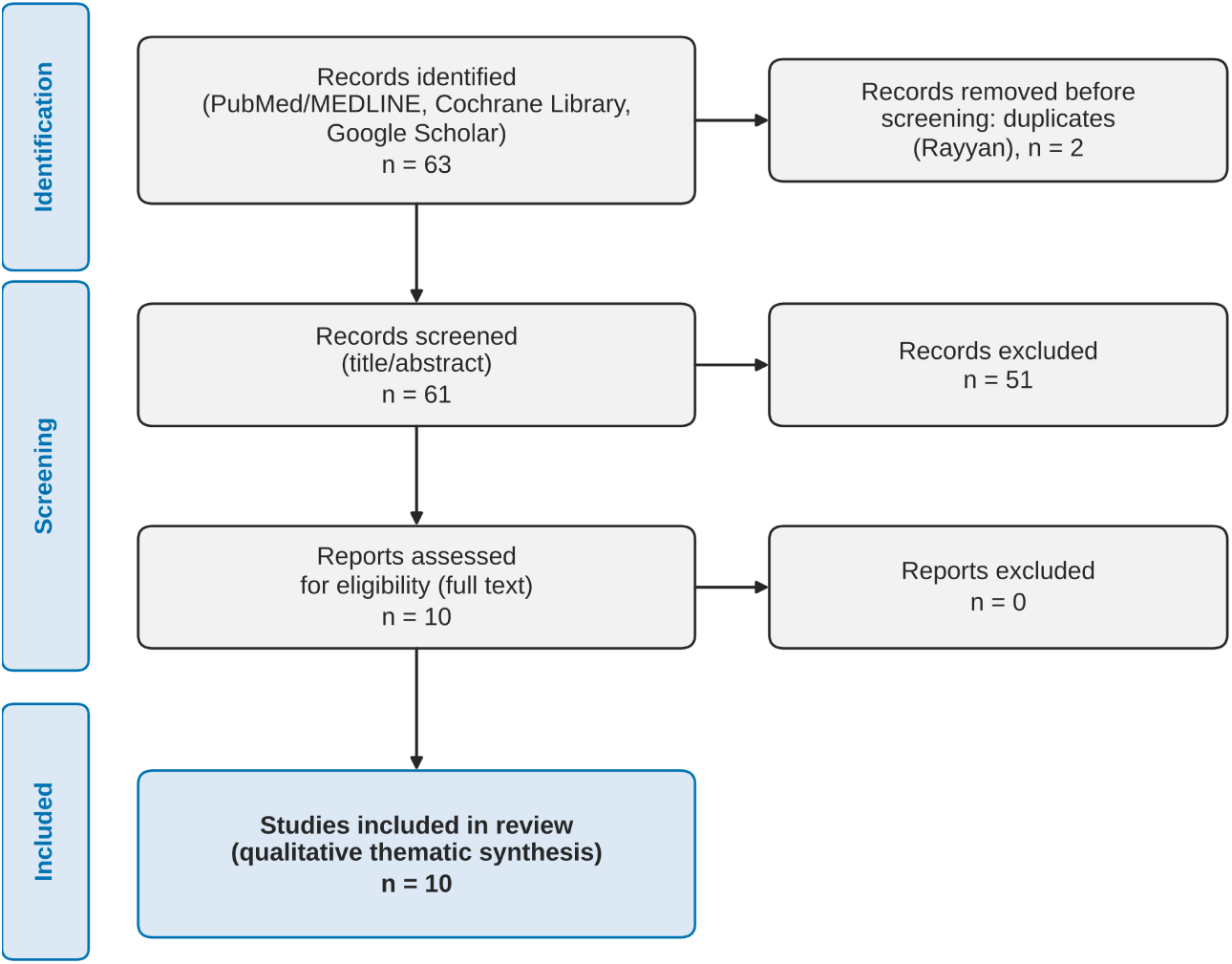
PRISMA 2020 flow diagram of study identification and selection. Records were deduplicated in Rayyan and screened independently. Exclusion criteria: adult-only cohorts, non-leukemic malignancies, publications without primary data and non-English-language publications. Adapted from the PRISMA 2020 template [24].

### 3.2 Characteristics of included studies

The ten included studies were published between 2016 and 2026 and originated from the USA (*n* = 4), China (*n* = 3), Belgium, Germany and Austria (*n* = 1 each) (Table 3; Figure 4) [11–13, 16, 20–23, 28, 29]. Seven analysed primary patient samples, two relied mainly on patient-derived xenografts and one used an engineered human induced pluripotent stem cell (hiPSC) model validated against patient samples. Cohort sizes ranged from 6 to 159 patients or samples. scRNA-seq was used in nine studies, scATAC-seq in four, CITE-seq and TCR/BCR sequencing in two each, and targeted single-cell DNA sequencing in one; the three largest datasets comprised 695,819, *>*540,000 and 136,671 cells.

**Figure 4:**
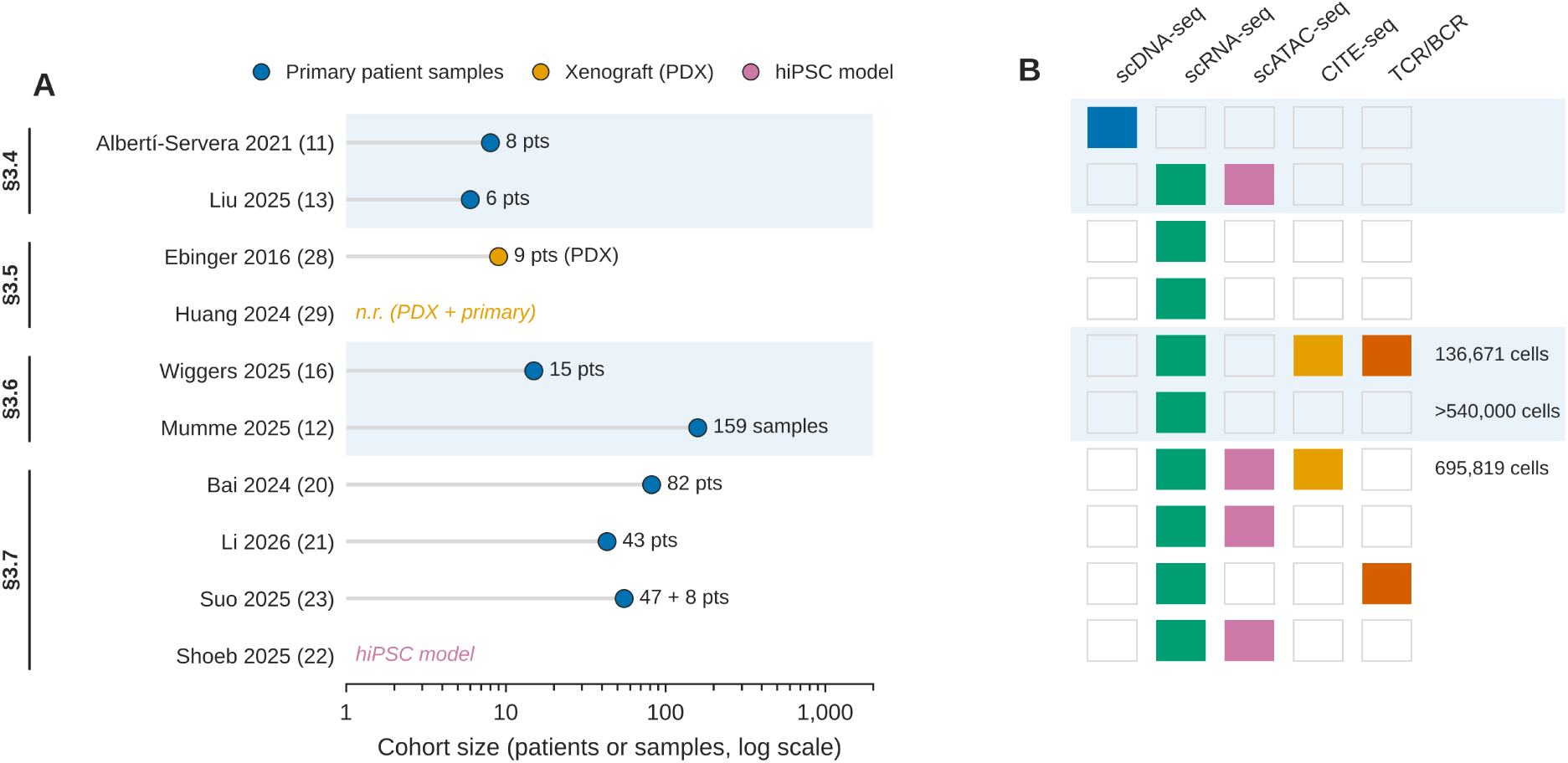
Characteristics of the ten included studies. (A) Cohort size (patients or samples, log scale), coloured by model system; studies are grouped by the Results section in which they are principally discussed (§3.4–3.7). n.r., not reported. (B) Single-cell modalities used by each study; values at right give total cells profiled where reported. Reference numbers in parentheses.

**Table 3:**
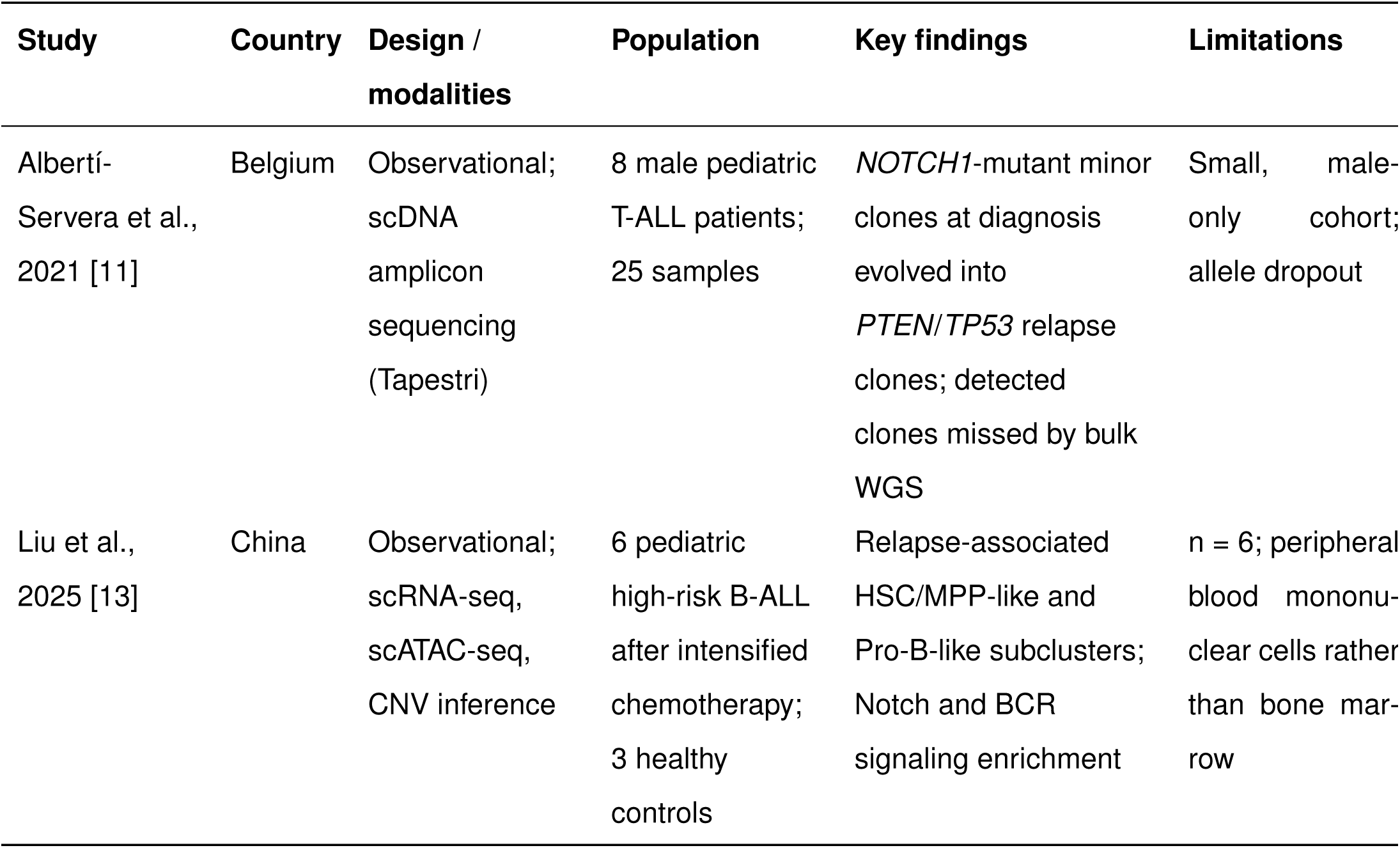
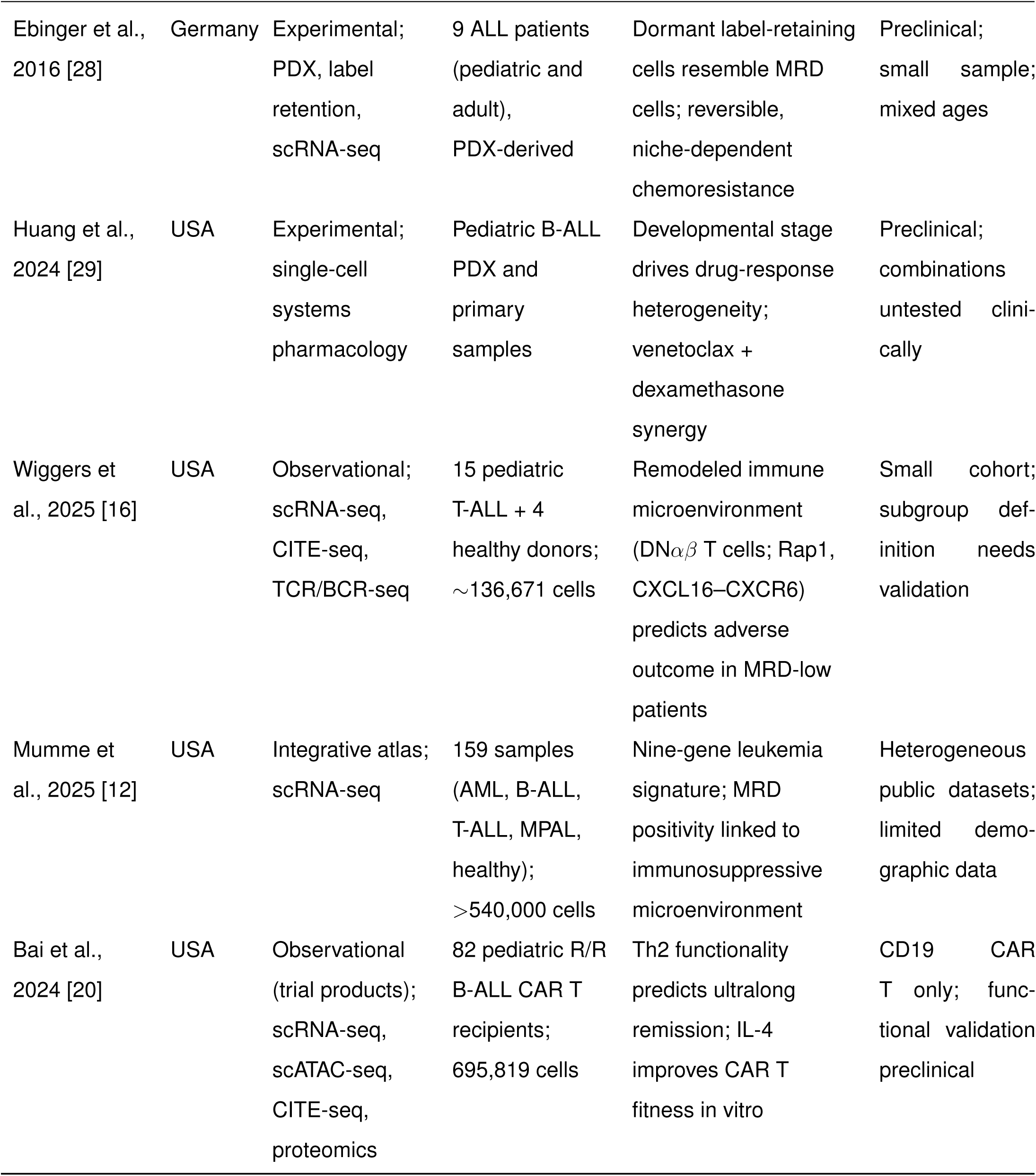
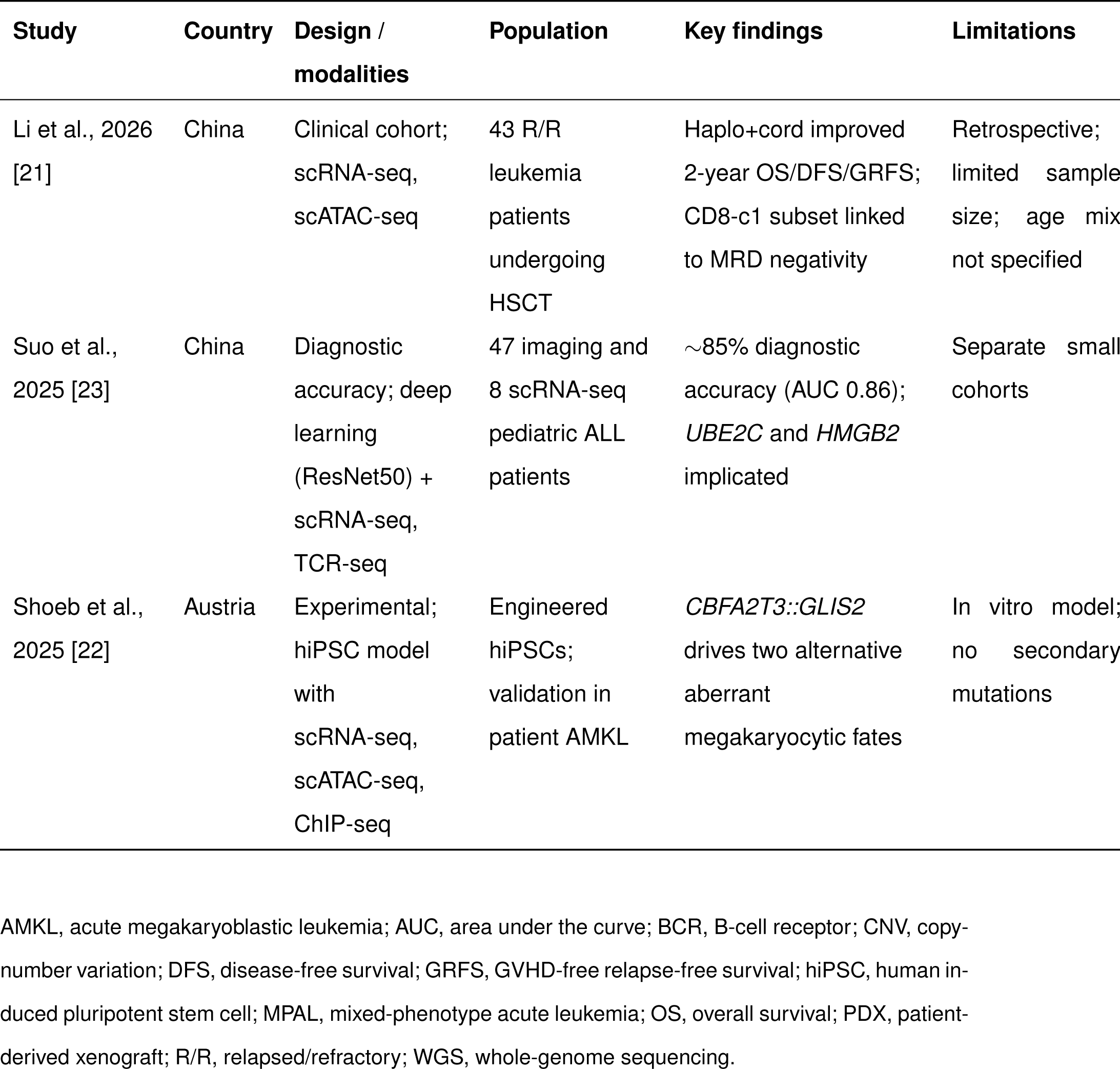
Characteristics and principal findings of the included studies.

| Study | Country | Design / modalities | Population | Key findings | Limitations |
| --- | --- | --- | --- | --- | --- |
| Alberti-Servera et al., 2021 [11] | Belgium | Observational; scDNA amplicon sequencing (Tapestri) | 8 male pediatric T-ALL patients; 25 samples | *NOTCH1*-mutant minor clones at diagnosis evolved into *PTEN/TP53* relapse clones; detected clones missed by bulk WGS | Small, male-only cohort; allele dropout |
| Liu et al., 2025 [13] | China | Observational; scRNA-seq, scATAC-seq, CNV inference | 6 pediatric high-risk B-ALL after intensified chemotherapy; 3 healthy controls | Relapse-associated HSC/MPP-like and Pro-B-like subclusters; Notch and BCR signaling enrichment | n = 6; peripheral blood mononuclear cells rather than bone marrow |
| Ebinger et al., 2016 [28] | Germany | Experimental; PDX, label retention, scRNA-seq | 9 ALL patients (pediatric and adult), PDX-derived | Dormant label-retaining cells resemble MRD cells; reversible, niche-dependent chemoresistance | Preclinical; small sample; mixed ages |
| Huang et al., 2024 [29] | USA | Experimental; single-cell systems pharmacology | Pediatric B-ALL PDX and primary samples | Developmental stage drives drug-response heterogeneity; venetoclax + dexamethasone synergy | Preclinical; combinations untested clinically |
| Wiggers et al., 2025 [16] | USA | Observational; scRNA-seq, CITE-seq, TCR/BCR-seq | 15 pediatric T-ALL + 4 healthy donors; ~136,671 cells | Remodeled immune microenvironment (DN$\alpha\beta$ T cells; Rap1, CXCL16–CXCR6) predicts adverse outcome in MRD-low patients | Small cohort; subgroup definition needs validation |
| Mumme et al., 2025 [12] | USA | Integrative atlas; scRNA-seq | 159 samples (AML, B-ALL, T-ALL, MPAL, healthy); >540,000 cells | Nine-gene leukemia signature; MRD positivity linked to immunosuppressive microenvironment | Heterogeneous public datasets; limited demographic data |
| Bai et al., 2024 [20] | USA | Observational (trial products); scRNA-seq, scATAC-seq, CITE-seq, proteomics | 82 pediatric R/R B-ALL CAR T recipients; 695,819 cells | Th2 functionality predicts ultralong remission; IL-4 improves CAR T fitness in vitro | CD19 CAR T only; functional validation preclinical |
| Li et al., 2026 [21] | China | Clinical cohort; scRNA-seq, scATAC-seq | 43 R/R leukemia patients undergoing HSCT | Haplo+cord improved 2-year OS/DFS/GRFS; CD8-c1 subset linked to MRD negativity | Retrospective; limited sample size; age mix not specified |
| Suo et al., 2025 [23] | China | Diagnostic accuracy; deep learning (ResNet50) + scRNA-seq, TCR-seq | 47 imaging and 8 scRNA-seq pediatric ALL patients | ~85% diagnostic accuracy (AUC 0.86); *UBE2C* and *HMGB2* implicated | Separate small cohorts |
| Shoeb et al., 2025 [22] | Austria | Experimental; hiPSC model with scRNA-seq, scATAC-seq, ChIP-seq | Engineered hiPSCs; validation in patient AMKL | *CBFA2T3::GLIS2* drives two alternative aberrant megakaryocytic fates | In vitro model; no secondary mutations |

AMKL, acute megakaryoblastic leukemia; AUC, area under the curve; BCR, B-cell receptor; CNV, copy-number variation; DFS, disease-free survival; GRFS, GVHD-free relapse-free survival; hiPSC, human induced pluripotent stem cell; MPAL, mixed-phenotype acute leukemia; OS, overall survival; PDX, patient-derived xenograft; R/R, relapsed/refractory; WGS, whole-genome sequencing.

### 3.3 Risk of bias in included studies

Of the seven studies assessed with ROBINS-I, four were at moderate and three at serious overall risk of bias (Figure 5; Supplementary Table S2). Selection of participants was the most problematic domain, rated serious in three studies because of very small, sex-restricted or high-risk-only cohorts [11, 13, 16]. Confounding was rated moderate in every study, reflecting uncontrolled differences in panel design, dataset preprocessing and conditioning regimens. Classification of interventions and deviations from intended interventions were consistently at low risk.

**Figure 5:**
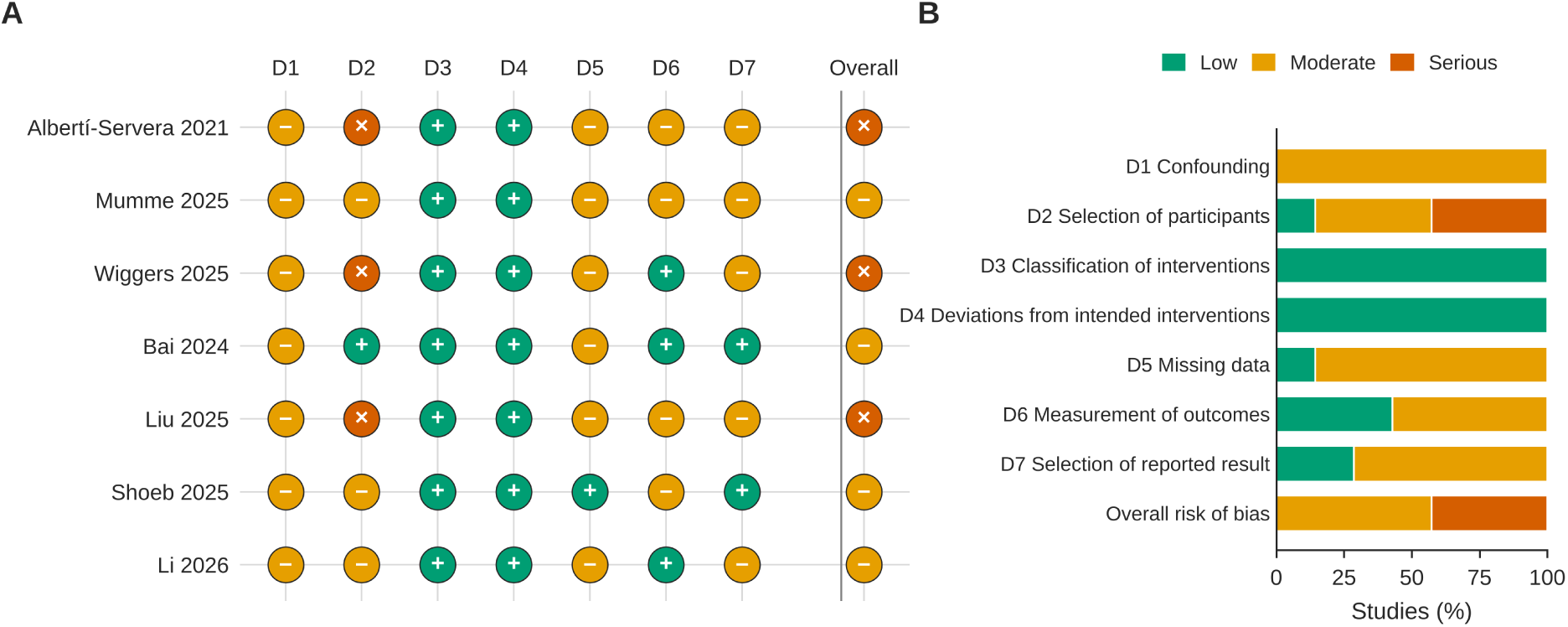
Risk of bias of included non-randomized studies (ROBINS-I). (A) Domain-level and overall judgements for each study; the overall judgement equals the most severe domain-level judgement. (B) Proportion of studies at each risk level per domain. D1, confounding; D2, selection of participants; D3, classification of interventions; D4, deviations from intended interventions; D5, missing data; D6, measurement of outcomes; D7, selection of the reported result. ROBINS-I was not applied to two xenograft-based experimental studies and one diagnostic-accuracy study (see Section 2.6).

### 3.4 Clonal evolution during treatment

The clearest demonstration of single-cell resolution of clonal evolution comes from Albertí-Servera et al., who applied single-cell DNA amplicon sequencing (Tapestri; 305 amplicons across 110 genes) to 25 bone marrow and peripheral blood samples from eight pediatric T-ALL patients [11]. Minor subclones present at diagnosis — in particular those carrying *NOTCH1* mutations — were not static bystanders but the direct precursors of relapse clones, frequently acquiring additional *PTEN* or *TP53* mutations (Figure 6A). Because single-cell genotyping resolves which mutations co-occur in the same cell, relapse-associated clones were detected in peripheral blood samples in which bulk whole-genome sequencing missed them entirely (Figure 6B).

**Figure 6:**
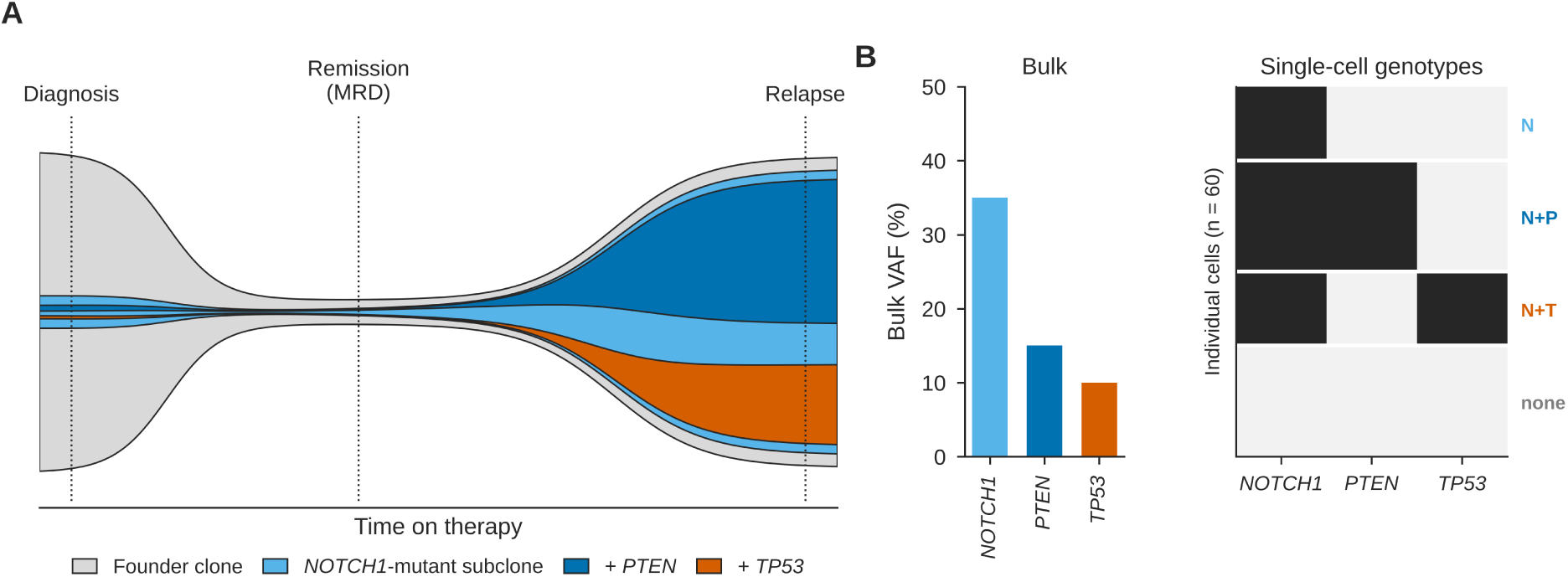
Clonal evolution from diagnosis to relapse resolved at single-cell resolution (schematic). (A) Fish plot of a T-ALL in which a minor *NOTCH1*-mutant subclone present at diagnosis survives therapy and, after acquiring *PTEN* or *TP53* mutations, dominates at relapse, as described by Albertí-Servera et al. [11]. Band height represents relative leukemic burden; nested bands denote descendant subclones. (B) Bulk variant allele frequencies (VAF; left) show which mutations are present, but only a cell *×* mutation matrix (right; black, mutant) shows that *PTEN* and *TP53* mutations arise in separate *NOTCH1*-mutant subclones. N, *NOTCH1*; P, *PTEN*; T, *TP53*; none, no tracked mutation. Illustrative data; proportions are not derived from patients.

Liu et al. integrated scRNA-seq and scATAC-seq in six pediatric patients with high-risk BALL after early intensified chemotherapy [13]. Non-remission samples showed expansion of hematopoietic stem/progenitor-like (HSC/MPP-like) and Pro-B-like subclusters, each with a distinct chromatin-accessibility signature: the HSC/MPP-like subcluster expressed *TCF4*, *EBF1*, *ERG* and *CRIM1* and was enriched for Notch signaling, whereas the Pro-B-like subcluster expressed *RPS29*, *B2M*, *RPL41* and *NEIL1* and was enriched for B-cell-receptor signaling. Clonal evolution under treatment pressure is therefore not limited to the acquisition of new mutations: shifts in the abundance and regulatory state of pre-existing developmental subpopulations can themselves constitute a route to treatment failure.

#### Contextual evidence

In adult AML, two large single-cell DNA sequencing cohorts established leukemia as a dynamic oligoclonal disease evolving along linear and branched trajectories, with higher clonal diversity at diagnosis associated with inferior survival [30, 31]. In pediatric BALL, single-cell transcriptomics has shown that each patient’s leukemia occupies a continuum of developmental states [32], and mtDNA-based lineage tracing offers a route to link clonal ancestry directly to transcriptional identity [9, 10].

### 3.5 Chemoresistant cell states

Ebinger et al. combined patient-derived xenografts of pediatric and adult ALL with proliferation-tracking dyes and single-cell and bulk RNA sequencing to identify a rare population of dormant, label-retaining cells [28]. These cells survived chemotherapy in vivo, retained leukemia-initiating potential and displayed reduced cell-cycle and DNA-replication gene expression with increased adhesion-pathway expression — a quiescence-like profile closely resembling that of clinical MRD cells. Dormancy and drug resistance were reversible once cells were removed from the bone marrow niche, implicating the microenvironment, not solely cell-intrinsic mutation, in chemoresistance.

Huang et al. applied scRNA-seq with systems-pharmacology profiling to pediatric B-ALL xenograft and primary samples [29]. The developmental stage of individual leukemic cells, rather than their mutational profile alone, was a major determinant of drug-response heterogeneity: sub-populations resistant to standard chemotherapy remained sensitive to targeted agents, and rationally selected combinations such as venetoclax with dexamethasone were synergistic against otherwise resistant clones. The Notch-and B-cell-receptor-enriched subclusters described by Liu et al. likewise point to potentially druggable pathways underlying treatment failure [13].

#### Contextual evidence

Relapse-fated B-ALL subclones that are intrinsically drug-tolerant before any treatment exposure have been isolated at diagnosis, distinguished by chromatin-remodeling, metabolic and stemness programs [18]. In *KMT2A*-rearranged infant ALL, glucocorticoid-resistant, quiescent stem-like cells detectable at diagnosis tracked with subsequent relapse [33]; in *ETV6-RUNX1*-positive ALL, chemoresistant populations showed cell-cycle arrest and loss of differentiation-associated transcription-factor activity that was pharmacologically targetable [34]; and scATAC-seq of therapy-resistant AML revealed a heterogeneous mixture of stem, progenitor and aberrantly differentiated cells rather than a single resistant state [14].

### 3.6 The immune microenvironment and MRD

Wiggers et al. profiled approximately 136,700 CD45^+^ immune cells from 15 pediatric T-ALL patients at diagnosis, during treatment and at remission, compared with healthy donors, using scRNA-seq, CITE-seq and TCR/BCR sequencing [16]. A subgroup comprising roughly 20–35% of patients displayed a remodeled immune microenvironment with enrichment of an unusual double-negative *αβ* T-cell population and non-classical monocytes, driven by leukemia-intrinsic Rap1 signaling and the CXCL16–CXCR6 axis. This signature was associated with adverse outcome specifically among patients classified as MRD-low — that is, immune profiling identified high-risk patients whom MRD assessment alone would have classified as favorable risk. Pharmacologic Rap1 inhibition sensitized leukemic cells to the BCL2/BCL-XL inhibitor navitoclax.

Mumme et al. integrated scRNA-seq data from 159 pediatric acute leukemia and healthy bone marrow samples (*>*540,000 cells) into a Pediatric Single-cell Cancer Atlas spanning AML, B-ALL, T-ALL and mixed-phenotype acute leukemia [12]. Beyond defining a nine-gene leukemia-enriched signature, the atlas showed that MRD positivity was consistently associated with depletion of naïve CD4^+^ and CD8^+^ T cells and M1 macrophages and enrichment of effector T cells and M2 macrophages — a shift toward an immunosuppressive microenvironment observed across leukemia subtypes. Together, these studies indicate that the immune microenvironment is mechanistically and prognostically intertwined with MRD, and that immune-aware profiling may add risk information beyond leukemic burden.

#### Contextual evidence

In pediatric AML, diagnostic samples from patients who later relapsed were enriched for exhausted T cells, whereas sustained remission was associated with inflammatory M1 macrophages [35]. In B-ALL, exhausted T cells up-regulating *TIGIT*, *PDCD1* and *LAG3* are present at diagnosis [36], and in relapsed T-ALL, exhausted CD8^+^ T and natural killer (NK) cells were identified, with JUN nominated as a driver of NK-cell exhaustion [37].

### 3.7 CAR T-cell persistence, post-transplant immune reconstitution and related applications

Bai et al. profiled 695,819 pre-infusion CD19-directed CAR T cells (CTL019) from 82 pediatric patients with relapsed or refractory B-ALL enrolled in two clinical trials, using scRNA-seq, scATAC-seq, CITE-seq and proteomics [20]. Elevated type 2 (Th2) functionality in the infusion product, regulated by GATA3 and STAT6, was strongly associated with ultralong remission (median duration of B-cell aplasia 8.4 years), and interleukin-4 (IL-4) supplementation during manufacturing alleviated CAR T-cell dysfunction in vitro — a manufacturing-actionable signature present before infusion that predicted response durability years later.

Li et al. used scRNA-seq and scATAC-seq to characterize immune reconstitution in 43 patients with relapsed or refractory leukemia undergoing haploidentical-plus-cord blood versus single cord blood HSCT [21]. The combined graft was associated with higher two-year overall survival (66.3% vs. 30.1%), disease-free survival (61.5% vs. 30.1%) and graft-versus-host-disease-free, relapse-free survival (54.2% vs. 20.3%) (Figure 7). Mechanistically, this benefit was linked to stronger type I interferon-mediated antitumor and antiviral immunity and a higher proportion of a cytotoxic, memory-marker-positive CD8^+^ T-cell subtype (CD8-c1; *CX3CR1*^+^*GNLY* ^+^) associated with MRD negativity and prolonged survival.

**Figure 7:**
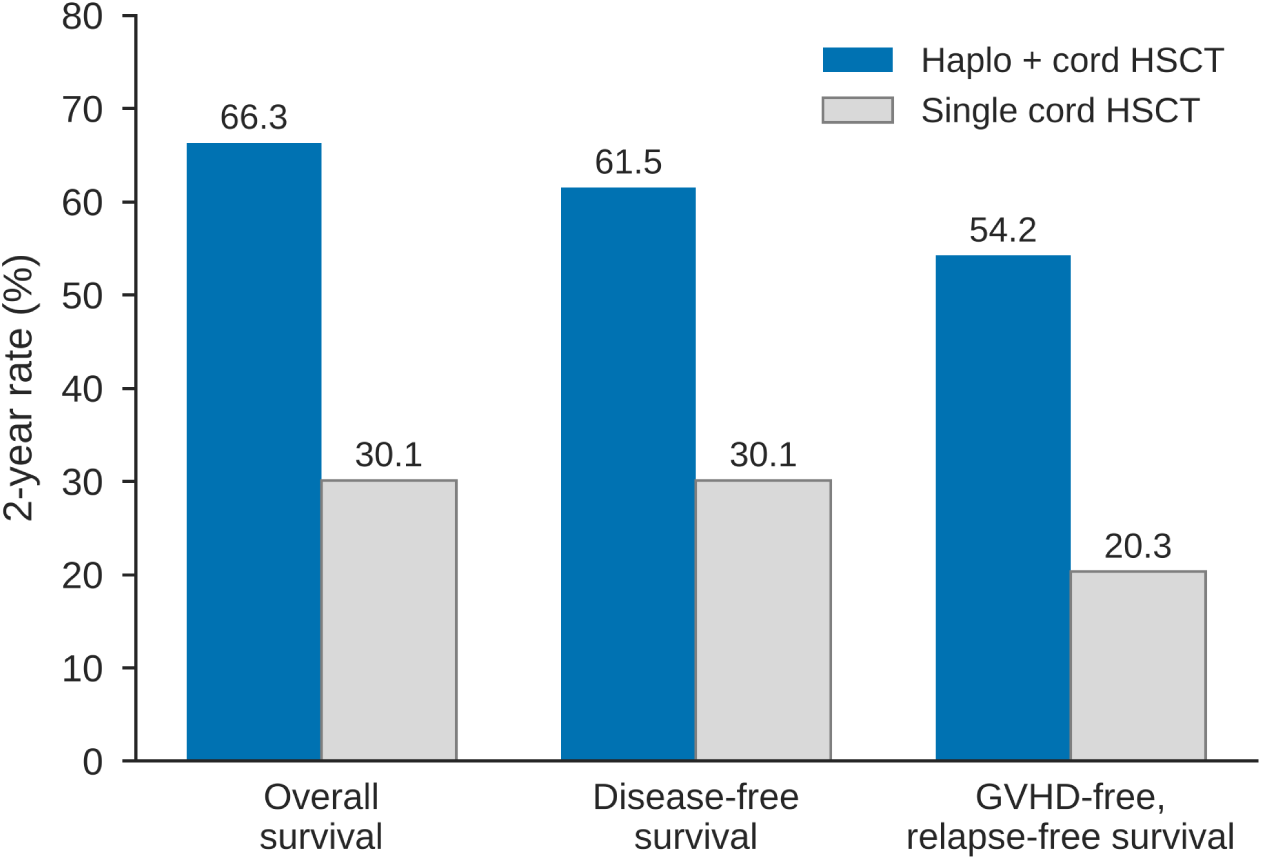
Two-year outcomes after haploidentical-plus-cord versus single cord blood transplantation. Overall, disease-free and graft-versus-host-disease-free, relapse-free survival in 43 patients with relapsed or refractory leukemia, as reported by Li et al. [21]. Confidence intervals were not available and are not shown.

Two further studies extend single-cell approaches to diagnostic and developmental questions. Suo et al. combined a ResNet50 deep-learning classifier of peripheral blood smear images with scRNA-seq and TCR sequencing, achieving approximately 85% diagnostic accuracy (area under the curve 0.86) for distinguishing ALL from normal cells and implicating *UBE2C* and *HMGB2* in leukemogenesis [23]. Shoeb et al. engineered hiPSCs to express *CBFA2T3::GLIS2*— the defining fusion of an aggressive pediatric acute megakaryoblastic leukemia subtype — and, using single-cell transcriptomic, chromatin-accessibility and chromatin-immunoprecipitation profiling, showed that the fusion drives two alternative aberrant megakaryocytic fates whose regulatory programs mirror those in primary patient samples [22].

#### Contextual evidence

In children treated on the CARPALL trial, long-lived CD19 CAR T cells converged on an exhausted-like memory state with a reproducible persistence signature [38], and a small study of infusion products found a relative deficit of Th2-functional and stem-like CAR T cells in patients who later relapsed with CD19-positive disease [39]. After allogeneic HSCT, the abundance of an immunoregulatory neutrophil-progenitor subset in the donor graft differed between patients who did and did not develop acute graft-versus-host disease [40], and infant *KMT2A*-rearranged B-ALL was shown to occupy a distinct early-lymphocyte-precursor-like state relative to fetal bone marrow [41]. A thematic cross-reference of the included studies is provided in Table 4.

**Table 4:**
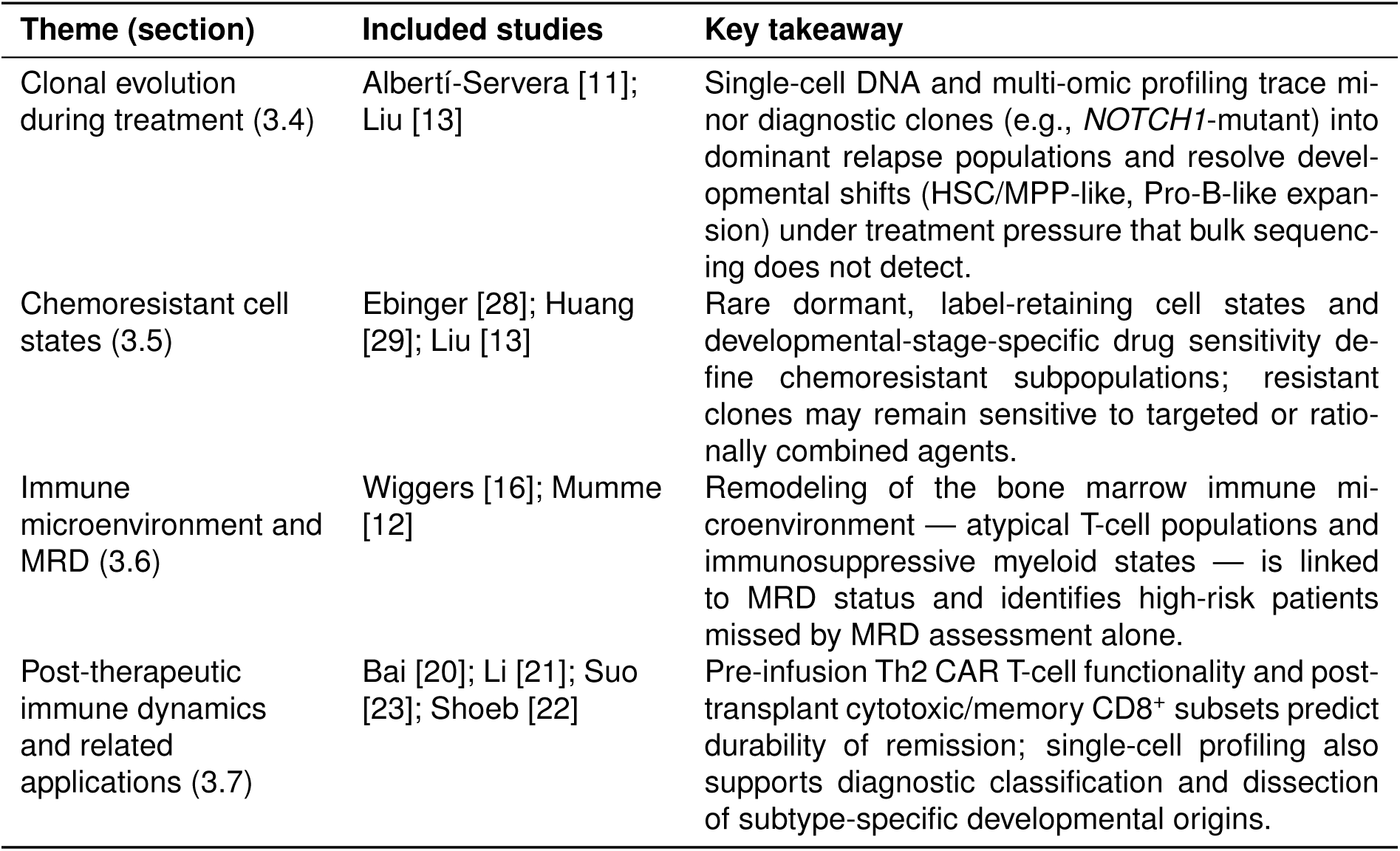
Key findings by biologic theme.

| Theme (section) | Included studies | Key takeaway |
| --- | --- | --- |
| Clonal evolution during treatment (3.4) | Albertí-Servera [11]; Liu [13] | Single-cell DNA and multi-omic profiling trace minor diagnostic clones (e.g., *NOTCH1*-mutant) into dominant relapse populations and resolve developmental shifts (HSC/MPP-like, Pro-B-like expansion) under treatment pressure that bulk sequencing does not detect. |
| Chemoresistant cell states (3.5) | Ebinger [28]; Huang [29]; Liu [13] | Rare dormant, label-retaining cell states and developmental-stage-specific drug sensitivity define chemoresistant subpopulations; resistant clones may remain sensitive to targeted or rationally combined agents. |
| Immune microenvironment and MRD (3.6) | Wiggers [16]; Mumme [12] | Remodeling of the bone marrow immune microenvironment — atypical T-cell populations and immunosuppressive myeloid states — is linked to MRD status and identifies high-risk patients missed by MRD assessment alone. |
| Post-therapeutic immune dynamics and related applications (3.7) | Bai [20]; Li [21]; Suo [23]; Shoeb [22] | Pre-infusion Th2 CAR T-cell functionality and post-transplant cytotoxic/memory CD8<sup>+</sup> subsets predict durability of remission; single-cell profiling also supports diagnostic classification and dissection of subtype-specific developmental origins. |

## 4 Discussion

### 4.1 Summary of evidence

Across ten included studies, single-cell sequencing consistently resolved four biologic processes that population-level clinical assays do not capture within a single test: the evolution of minor diagnostic subclones into dominant relapse clones; the existence of rare dormant or developmentally distinct cell states that tolerate chemotherapy; immune-microenvironment remodeling that tracks with MRD and identifies adverse outcome among MRD-low patients; and immune cell states at CAR T-cell infusion or after HSCT that predict durability of remission. These findings are mutually reinforcing and are corroborated by contextual single-cell studies outside the formal synthesis. Their certainty is nonetheless limited by small cohorts, moderate-to-serious risk of bias and the absence of prospective validation.

### 4.2 Potential clinical applications

The synthesized findings point toward four plausible, though not yet validated, clinical applications (Figure 8).

**Figure 8:**
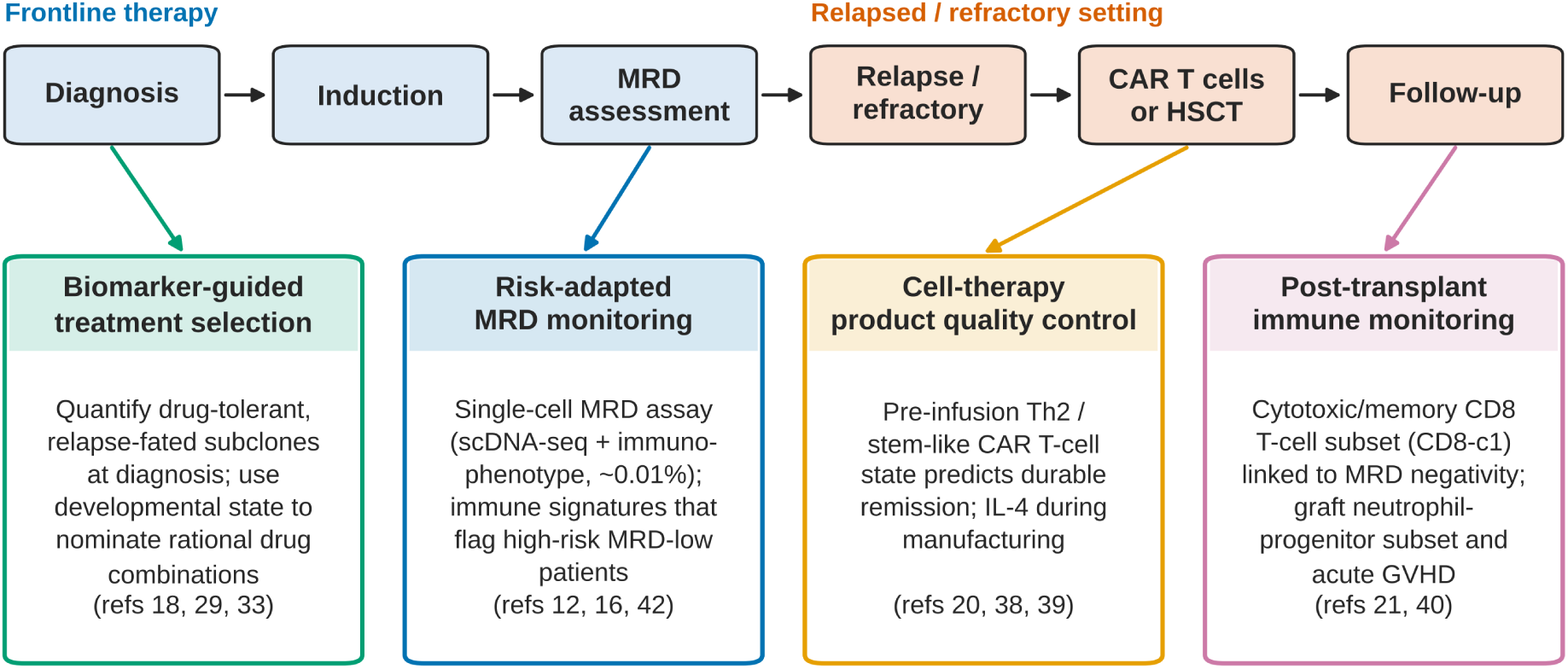
Potential points of clinical application of single-cell sequencing along the pediatric acute leukemia care pathway. Each application corresponds to a paragraph of Section 4.2 and lists the supporting references. None of these applications is yet clinically validated; prospective, harmonized studies are required (Section 4.3).

#### Risk-adapted MRD monitoring

Immune signatures such as those described by Wiggers et al. and Mumme et al. could refine MRD-based risk stratification, particularly for MRD-low patients whose immune microenvironment predicts adverse outcome [12, 16]. A dedicated single-cell MRD assay is the most direct near-term opportunity: flow cytometry can miss cells with ambiguous immunophenotypes, whereas bulk NGS cannot reliably distinguish residual leukemia from clonal hematopoiesis. An assay combining progenitor enrichment with integrated single-cell DNA sequencing and immunophenotyping has achieved an estimated sensitivity on the order of 0.01% in AML [42]; in pediatrics, such an approach could be most valuable in AML and lineage-ambiguous leukemias, where flow-and PCR-based MRD are least reliable.

#### Biomarker-guided treatment selection

The observation that developmental stage and chromatin state, rather than mutational profile alone, predict drug sensitivity [13, 29], together with evidence that relapse-fated drug-tolerant subclones are already present at diagnosis [18, 33], suggests that single-cell profiling at diagnosis could flag patients for earlier escalation to targeted therapy or immunotherapy and nominate rational drug combinations. This remains exploratory and would require demonstration of improved outcomes in prospective trials.

#### Cell-therapy manufacturing and quality control

The association between pre-infusion Th2 functionality and remission durability [20], reinforced by independent persistence signatures [38, 39], is among the most immediately actionable findings: it identifies a measurable manufacturing-stage biomarker and a specific intervention (IL-4 supplementation) that could be tested in early-phase trials, with particular relevance to pediatric B-ALL, in which loss of CAR T-cell persistence is a recognized mechanism of relapse.

#### Post-transplant immune monitoring

The CD8-c1 cytotoxic memory T-cell subtype associated with graft source and favorable outcome [21], and the graft neutrophil-progenitor subset associated with acute graft-versus-host disease [40], raise the possibility of single-cell immune monitoring after HSCT to identify patients at higher risk of relapse or graft-versus-host disease before these become clinically apparent.

### 4.3 Limitations of the evidence

Most included studies used small, single-institution cohorts (6–159 patients or samples), limiting statistical power and generalizability. Platforms, sequencing depth and computational pipelines differed substantially between studies, complicating direct comparison. Three studies relied on xenograft or hiPSC models that, while mechanistically informative, may not recapitulate the in vivo leukemic microenvironment. Nearly all findings derive from retrospective analyses; no study linked a single-cell biomarker prospectively to outcome and treatment decisions. More broadly, single-cell assays remain costlier, slower and more computationally demanding than flow cytometry or PCR; no standardized clinical protocol or analytic pipeline exists; and, as laboratory-developed multi-omic tests, single-cell MRD and related assays will require formal analytical validation and a defined regulatory pathway.

### 4.4 Limitations of the review process

This review has several limitations. First, the search yielded a small number of records, and several pediatric single-cell studies that would plausibly meet the eligibility criteria — including studies of infant *KMT2A*-rearranged ALL, *ETV6-RUNX1*-positive ALL, pediatric AML and CAR T-cell persistence [33–35, 38, 41] — were not retrieved by the structured search; these are reported as contextual evidence, but their absence from the formal synthesis indicates limited search sensitivity. Second, Google Scholar searches are not fully reproducible. Third, the protocol was not prospectively registered. Fourth, ROBINS-I was developed for non-randomized studies of interventions and was adapted here to observational and model-based designs; it was not applied to three studies. Fifth, certainty of evidence was not graded and reporting biases were not formally assessed. Finally, restriction to English-language publications may have excluded relevant work.

### 4.5 Implications for research

Priorities include multi-center, prospective cohorts with harmonized sample processing and analytic pipelines; prespecified evaluation of single-cell-derived biomarkers as independent predictors alongside standard MRD; early-phase trials testing manufacturing interventions informed by pre-infusion CAR T-cell states; and analytical validation of single-cell MRD assays in pediatric AML and lineage-ambiguous leukemias. Shared reference atlases, such as pediatric single-cell atlases [12], will be essential for cross-study comparability.

## 5 Conclusions

Single-cell sequencing has generated mechanistic insight into clonal evolution, chemoresistance, immune dysregulation and post-therapeutic immune dynamics in pediatric acute leukemia that is not obtainable from conventional population-level assays, and it complements rather than replaces clinically validated MRD testing. The evidence remains drawn predominantly from small, retrospective and often preclinical studies at moderate-to-serious risk of bias. Larger, prospective and harmonized studies are required before single-cell-derived biomarkers can be incorporated into risk stratification or treatment algorithms, but the biologic rationale for doing so is increasingly well supported.

## Declarations

### Author contributions

J.M.: conceptualization, literature search, study selection, data extraction, risk-of-bias assessment, visualization and writing.

### Data and code availability

All extracted data are reported in the tables and supplementary material. The Python code used to generate the figures is available as Supplementary File 1 and from the corresponding author on request.

### Use of generative AI

Generative AI (Claude, Anthropic) was used to assist with restructuring and language editing of the manuscript and to write the Python (matplotlib) code used to produce the figures. All figure values are taken from the included studies as reported; the author reviewed and verified all content and takes full responsibility for the manuscript.

## Acknowledgements.

None.

## Supplementary Material

**Supplementary Table S1.**
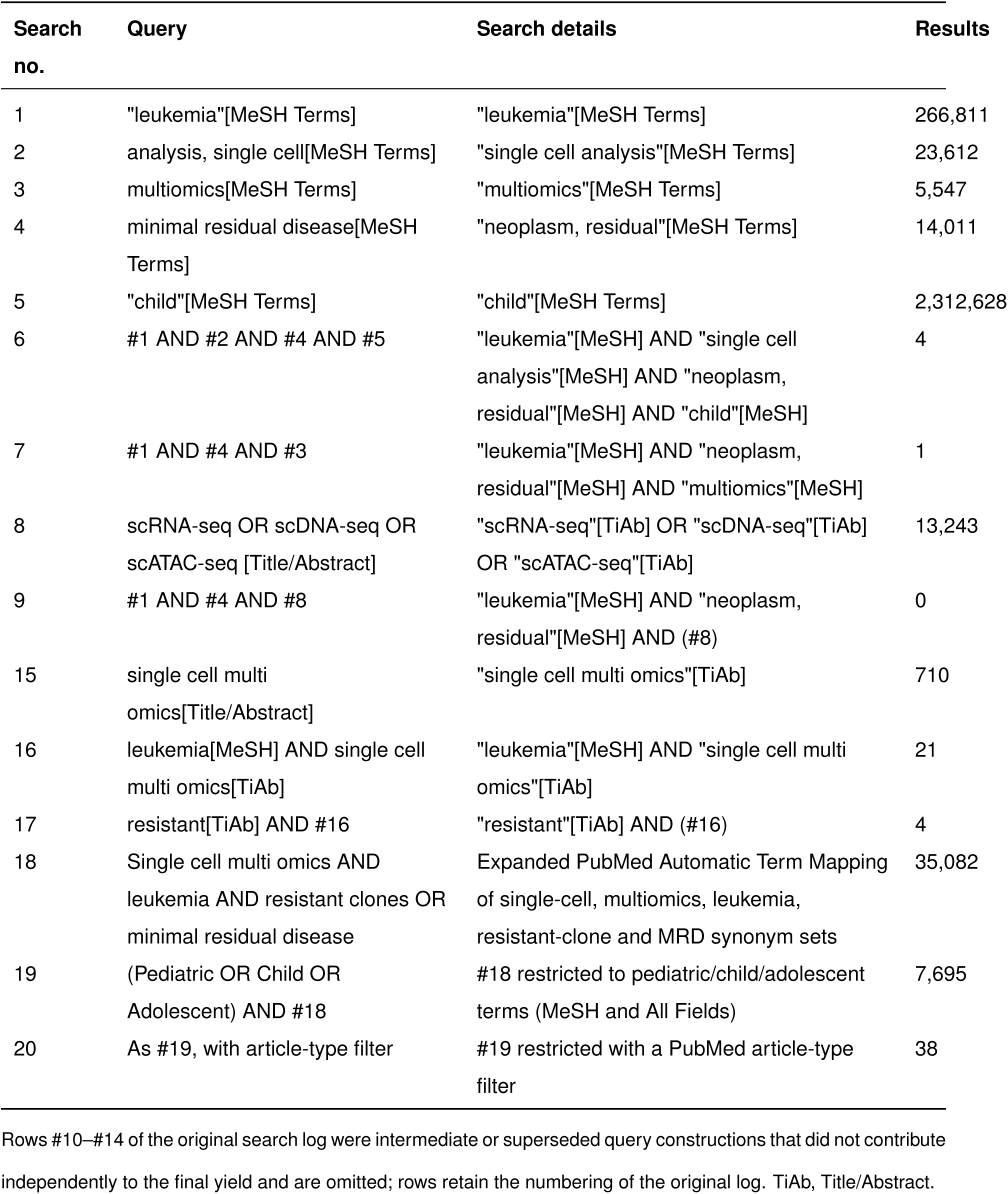
Full PubMed search output. Rows #10–#14 of the original search log were intermediate or superseded query constructions that did not contribute independently to the final yield and are omitted; rows retain the numbering of the original log. TiAb, Title/Abstract.

| Search no. | Query | Search details | Results |
| --- | --- | --- | --- |
| 1 | "leukemia"[MeSH Terms] | "leukemia"[MeSH Terms] | 266,811 |
| 2 | analysis, single cell[MeSH Terms] | "single cell analysis"[MeSH Terms] | 23,612 |
| 3 | multiomics[MeSH Terms] | "multiomics"[MeSH Terms] | 5,547 |
| 4 | minimal residual disease[MeSH Terms] | "neoplasm, residual"[MeSH Terms] | 14,011 |
| 5 | "child"[MeSH Terms] | "child"[MeSH Terms] | 2,312,628 |
| 6 | #1 AND #2 AND #4 AND #5 | "leukemia"[MeSH] AND "single cell analysis"[MeSH] AND "neoplasm, residual"[MeSH] AND "child"[MeSH] | 4 |
| 7 | #1 AND #4 AND #3 | "leukemia"[MeSH] AND "neoplasm, residual"[MeSH] AND "multiomics"[MeSH] | 1 |
| 8 | scRNA-seq OR scDNA-seq OR scATAC-seq [Title/Abstract] | "scRNA-seq"[TiAb] OR "scDNA-seq"[TiAb] OR "scATAC-seq"[TiAb] | 13,243 |
| 9 | #1 AND #4 AND #8 | "leukemia"[MeSH] AND "neoplasm, residual"[MeSH] AND (#8) | 0 |
| 15 | single cell multi omics[Title/Abstract] | "single cell multi omics"[TiAb] | 710 |
| 16 | leukemia[MeSH] AND single cell multi omics[TiAb] | "leukemia"[MeSH] AND "single cell multi omics"[TiAb] | 21 |
| 17 | resistant[TiAb] AND #16 | "resistant"[TiAb] AND (#16) | 4 |
| 18 | Single cell multi omics AND leukemia AND resistant clones OR minimal residual disease | Expanded PubMed Automatic Term Mapping of single-cell, multiomics, leukemia, resistant-clone and MRD synonym sets | 35,082 |
| 19 | (Pediatric OR Child OR Adolescent) AND #18 | #18 restricted to pediatric/child/adolescent terms (MeSH and All Fields) | 7,695 |
| 20 | As #19, with article-type filter | #19 restricted with a PubMed article-type filter | 38 |

**Supplementary Table S2.**
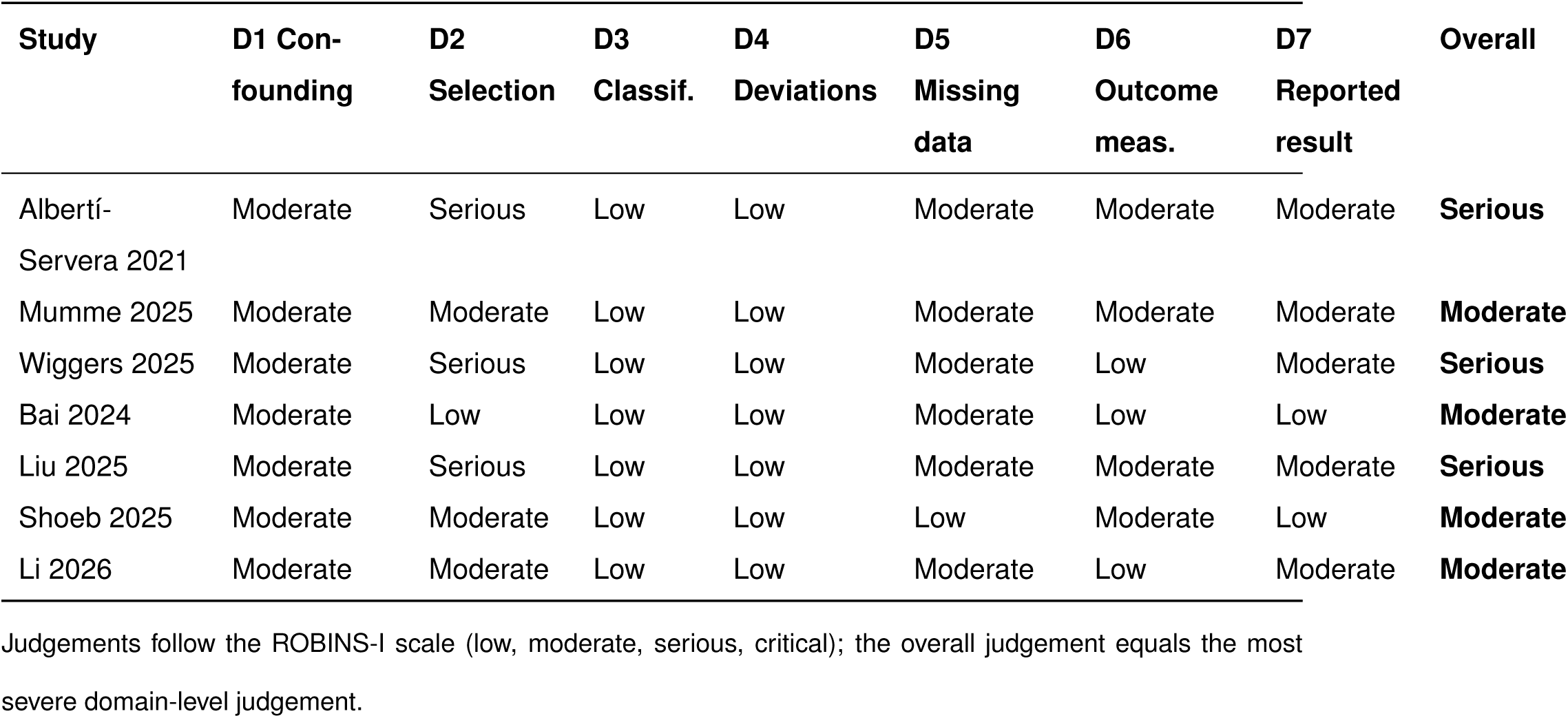
ROBINS-I domain-level judgements and rationale. Judgements follow the ROBINS-I scale (low, moderate, serious, critical); the overall judgement equals the most severe domain-level judgement.

**Supplementary Table S3.**
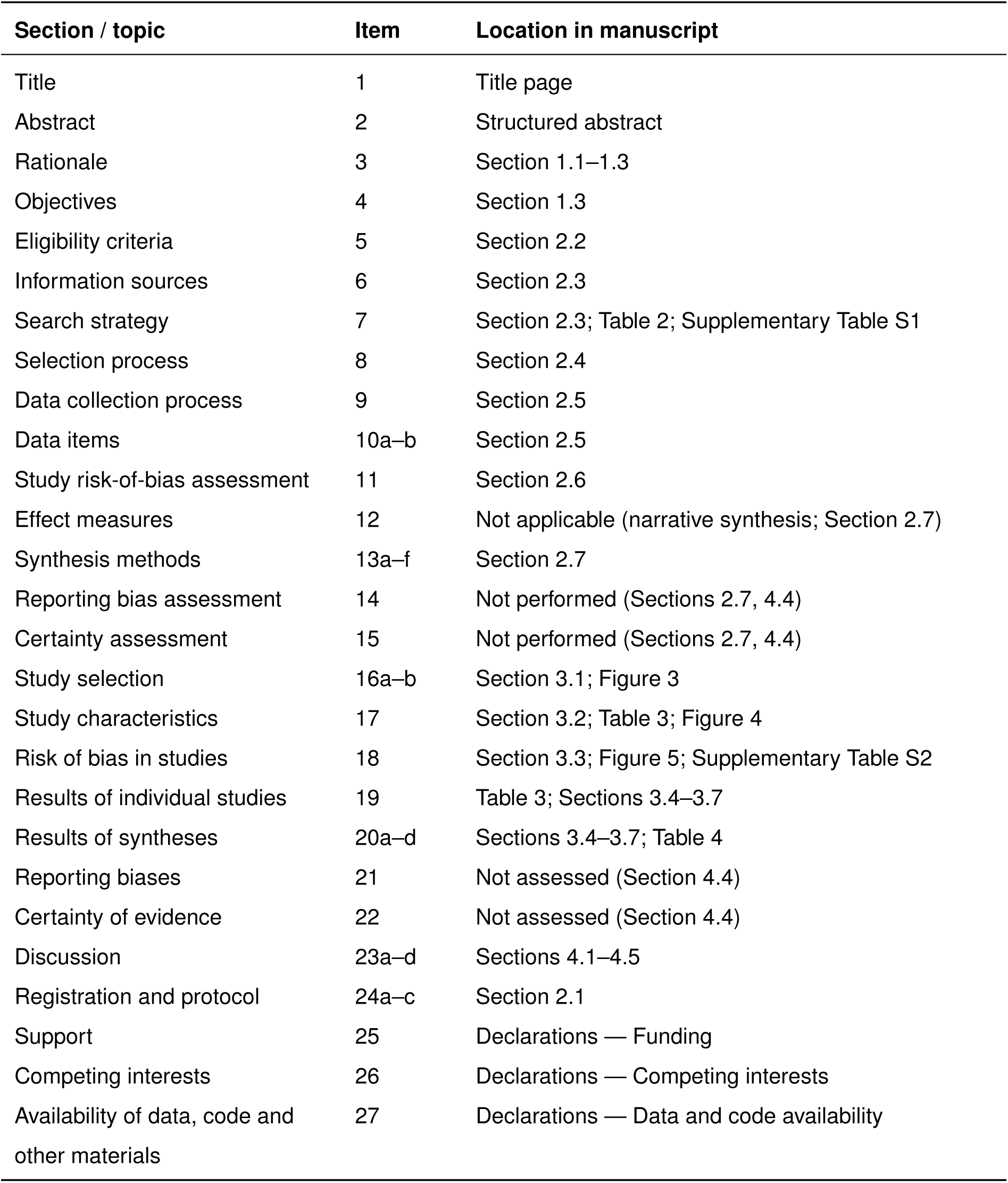
PRISMA 2020 checklist.

